# Red-purple Andean potato polyphenols have an anti-neuroblastoma effect *in vitro* via apoptosis through mitochondria

**DOI:** 10.1101/2023.02.01.526690

**Authors:** María Ximena Silveyra, Adriana Balbina Andreu

## Abstract

Andean potatoes (*Solanum tuberosum* ssp. *andigena*) are a good source of dietary polyphenols, such as phenolic acid and flavonoids. These polyphenols have several beneficial effects on human health due to their antioxidant properties. Previously, we demonstrated that polyphenol extracts from Andean potato tubers exerted a concentration-dependent cytotoxic effect in human neuroblastoma cells. However, the mechanisms involved in this cytotoxic activity were not explored. Here we show that Santa María tuber’s polyphenols activated a programmed cell death by caspase-independent apoptosis. We found that polyphenols induced cell and nucleus morphology changes and slightly affected the cell cycle. Furthermore, the polyphenols altered the neuroblastoma cells’ homeostasis redox and mitochondrial function, increasing the levels of apoptotic cells. Finally, we showed that neither Bcl-2 nor caspase-3 was involved in this mechanism of death. Our results confirmed that Santa María tuber’s polyphenols are bioactive compounds with mitochondria as a target and contribute to revalorizing Andean potatoes as a functional food. These findings demonstrated that they would be a good source of anti-tumor compounds that would induce tumor cell death even in apoptotic-resistant tumors, opening new therapeutic avenues.

## Introduction

Andean potatoes (*Solanum tuberosum* ssp. *andigena*) are native to South America and present high genetic biodiversity. These tubers are food staples for the Andean population, and those varieties with high nutritional properties were selected and cultivated over the years^1^. Potato is highly nutritious due to its high starch, vitamins, minerals, and fibers. In particular, red and purple Andean potatoes have significant amounts of antioxidant polyphenols such as phenolic acid, flavanols, flavanones, and anthocyanins ^2^.

*Solanum tuberosum* ssp. *andigena* var. Santa María (Accession CL 658 Santa María) is one of the 48 potato varieties collected and preserved in the Active Germplasm Bank of INTA-EEA Balcarce (BAL), Buenos Aires, Argentina^3^. Santa María’s tuber is small and elongated, and it has intense red-purple skin and purple flesh^1^. In the flesh of this variety, the major phenolic compounds are anthocyanidins (65.4%), while phenolic acids represent 31%. The abundant compounds are pelargonidin and chlorogenic acid, followed by peonidin, malvidin, and caffeic acid^2^.

Polyphenols are associated with specific beneficial effects on human health, acting as antioxidant compounds capable of preventing or delaying molecular damage caused by reactive oxygen species (ROS). In the first instance, incorporating foods rich in antioxidant polyphenols into the diet would protect the body against oxidative stress that causes many chronic diseases, such as cancer^4^. Furthermore, these compounds can act indirectly by activating endogenous defense systems and intervening in intracellular signaling pathways, conferring other beneficial attributes^5,6^. In addition, recent investigations have demonstrated that polyphenols act on mitochondria, the organelle responsible for producing energy and intracellular ROS^7,8^.

The polyphenols present in the potato have gained interest in human health due to their antioxidant properties^9–11^. Extensive evidence shows that potato has beneficial properties in cell cultures and animal models, with positive effects against oxidative stress, hypercholesterolemia, inflammation, obesity, diabetes, and cancer^12^. Thus, potato polyphenols are potential anti-tumoral compounds for several cancers. For example, potato polyphenols extracts showed antiproliferative activity in the human colon, breast, liver, and leukemia cells^13–18^. In particular, anthocyanins-fraction from colored potatoes induced cytotoxicity and apoptosis in prostate cancer cells^19^. Even purple-fleshed potatoes keep pro-apoptotic properties after baking, demonstrating that Andean potatoes could be a functional food contributing to human health^20^.

Our group recently demonstrated that Santa María polyphenols have cytotoxic activity mediated by apoptosis and autophagy in human hepatocarcinoma cells^21^. We have also shown that polyphenol extracts from Andean potato tubers exerted a concentration-dependent cytotoxic effect in human neuroblastoma cells^22^. However, the mechanisms involved in this preventive activity were not explored. We wondered if this effect occurs through a controlled cell death mechanism or massive necrosis capable of triggering an unwanted inflammatory response. This work investigates the death mechanism involved in the anti-neuroblastoma activity by the flesh tuber’s polyphenols from the Andean potato variety Santa María. We performed microscopy and biochemical experiments on an *in vitro* neuroblastoma cell model. Our results showed that the Santa María tuber’s polyphenols altered the neuroblastoma cells’ homeostasis redox and mitochondrial function, activating a programmed cell death by caspase-independent apoptosis. This effect of Santa María’s polyphenols provides evidence for using Andean potatoes as a functional food to be included in the diet to promote health and prevent diseases.

## Materials and methods

### Biological Materials

The Native Andean potato variety Santa María was selected based on previous work from our laboratory^2,22^. This variety was grown in the Northwest of Argentina in a field located in the Yavi Department (22° 6’ 4” S, 65° 35’ 44” O, 3377 MAMSL), Jujuy, Argentina. Potatoes were harvested and transported to the laboratory, where they were washed with tap water and kept at 4°C until processing. Each potato was peeled and cut into 1 x 1 cm pieces at room temperature, frozen into liquid nitrogen, and kept at -80°C until analysis. Then, the flesh pieces were freeze-dried for three days using a lyophilizer (Model L-A-B3, Rificor).

### Preparation of polyphenol extracts from Andean potatoes

Lyophilized flesh was finely powdered, and 1 g of powder was extracted with 20 mL of 90% methanol (v/v) (HPLC-grade, Baker) at 4°C in darkness overnight. Then, the homogenate was centrifugated at 4000 *g* for 20 min, and the supernatant was filtered using a muslin mesh. The extract was concentrated at 43°C using a speed vacuum concentrator (RVC 2-18 CD plus, Martin Christ) and resuspended in 1 mL of 5% DMSO (v/v). We named the polyphenol extracts PFE (<u>P</u>otato <u>F</u>lesh <u>E</u>xtract) to simplify.

### Total phenolic content

We determined the total phenolic content according to a modified Folin-Ciocalteu method. Briefly, the extract was diluted in 20 μL of water and loaded on a 96-well plate. To each well, 7% Folin-Ciocalteu reagent (v/v) was mixed with the sample and incubated at room temperature in darkness for 5 min. Then, 500 mM sodium carbonate was added and allowed for color development. Finally, we measured absorbance at 725 nm with a microplate spectrophotometer (Epoch BioteK). We extrapolated the data using a standard curve with chlorogenic acid (CGA), and we expressed the total phenolic acid content as μg CGA equivalents per mL (μg/mL).

### Cell culture and treatments

The cell line SH-SY5Y (ATCC^®^ Catalog No. CRL-2266™) was maintained in DMEM-F12 medium (Invitrogen, ThermoFisher Scientific) supplemented with 10% fetal bovine serum (v/v) (Natocor), penicillin (100 U/mL), streptomycin (100 μg/mL), and amphotericin B (0.25 μg/mL) (Invitrogen, ThermoFisher Scientific) at 37°C in a humidified atmosphere containing 95% air and 5% CO2. For the treatments, cells were seeded in 24-well plates at 50 x 10^4^ cells/well for 48 hr and subsequently incubated in reduced serum medium in the presence of a vehicle (less than 0.1% DMSO) or 400 μg/mL of PFE. In previous experiments, we determined this concentration as the CC_50_. After treatment, we observed the morphology of the cells by contrast microscopy (Olympus CKX41). We captured the images using the software Q-Capture Pro7.To investigate how the cells dye, the treatment time was different for each experiment depending on the mechanism evaluated. In the case of ROS levels and mitochondrial membrane potential, the incubation time was shorter at the same time as the measurement.

### Nuclei morphology

Since the DNA would be a final target in apoptosis, we treated the cells for 48 hr and fixed them with 4% formaldehyde at room temperature for 10 min. Then, the cells were washed with 1X PBS, resuspended with 0.2 mL of 1X PBS, and incubated with 0.2 µg/mL of DAPI at room temperature in darkness for 30 min. Finally, we spun down the cells and loaded them onto a microscope slide with a mounting solution (70% glycerol). Cells were analyzed immediately by fluorescence microscopy with a 100x oil immersion objective (Nikon Eclipse E200). We set the same time of exposure to compare control and treated cells. We measured the nuclei diameter and the fluorescence intensity of at least 200 cells using ImageJ software (version 1.53k, National Institutes of Health).

### Cell cycle and sub-G1 analysis

After 48 hr treatment, the cells were fixed with dropwise 70% cold-ethanol at 4°C for at least 2 hr, centrifuged, and resuspended in 1X PBS. We added 100 µg/mL Ribonuclease and 50 µg/ml propidium iodide (PI, SIGMA Aldrich) and incubated at room temperature in darkness for 30 min. We analyzed 50000 cells using a flow cytometer (Partec GmbH, Lab Systems SA), setting the FL-3 channel for PI on a linear scale. The coefficient of variation of the calibration beads peak on the FL-3-channel was less than 6%. The histogram of DNA fluorescence represents the G1, S, and G2/M cell cycle phases.

### Measurement of cell apoptosis

The cells treated for 24 hr were washed with 1X Binding Buffer and resuspended in 50 µL of the same buffer. We incubated with 5 µL of Annexin V-conjugated FITC (MACS Miltenyi Biotec, Lab Systems SA) at room temperature in darkness for 20 min. Then, we completed the volume to 1 mL final with 1X Binding Buffer and filtered the sample through a 50-µm nylon filter into a polypropylene tube. Finally, we added 1 µg/mL PI immediately before analysis by flow cytometry. We analyzed 50000 cells using the FL-1 channel for Annexin V-FITC and the FL-3 channel for PI on a logarithmic scale.

### Determination of intracellular reactive oxygen species (ROS)

We replaced the cell growth media with a reduced serum medium containing 20 µM of H_2_DCFDA (Molecular Probes, Invitrogen) and incubated it at 37°C in darkness for 1 hr. After loading the probe, we removed it and treated cells with the CC_50_ of PFE. We added 200 µM of H_2_O_2_ as a positive control. Immediately, we read the fluorescence intensity every 60 min for 4 hr using a fluorometer (Ex 488/Em 525, Fluoroskan Ascent, ThermoFisher Scientific). The fluorescence intensity was measured in 40 different points distributed in the entire area of the well with 50 x 10^4^ cells.

### Measurement of the mitochondrial membrane potential

To measure the kinetics of the mitochondrial membrane potential, we loaded 1 µg/µL Rhodamine 123 (ThermoFisher Scientific) in a reduced serum medium in darkness at 37°C for 10 min. Then, we treated the cells with the CC_50_ of PFE for 60 min, and we read the fluorescence intensity using a fluorometer (Ex 488/Em 525) every 15 min. On another approach, we first treated the cells for 3 hr, loaded the dye, and read the fluorescence. In both, the fluorescence intensity was measured in 40 different points distributed in the entire area of the well with 50 x 10^4^ cells.

### DNA fragmentation

As DAPI staining, we lysed the cells with 0.25 mL of Lysis Buffer (50 mM Tris-HCl pH 8, 10 mM EDTA pH 8, 100 mM NaCl, 0.5% SDS) after 48 hr treatment. Then, we incubated each sample with 50 µg/mL RNAse at 37°C for 30 min and with 0.1 mg/mL Proteinase K at 56°C for 1-2 hr. We precipitated the genomic DNA with 1/10 volume of 3 M sodium acetate pH 5.2 and two volumes of 100% ethanol and incubated at -20°C overnight. We washed the DNA with 70% and 100% ethanol, and the pellet was air-dried (5 min) and resuspended in 20 µL of TE pH 8. The DNA was separated electrophoretically on a 1% agarose gel (TBE 1X) containing 0.5 µg/mL Ethidium Bromide. We loaded 5 µL of qLadder 100 pair bases precision (PB-L Productos Bio-Lógicos®) as a molecular weight marker. The run was at 90V for 1 hr, and we visualized the DNA by ultraviolet transillumination.

### Western blotting

For protein extraction, cells were lysed after 24 hr of treatment on ice in HEPES buffer (50 mM Hepes, pH 7.4, 150 mM NaCl, 10% Glycerol, 1% Triton X-100, 1 mM EDTA, 1.5 mM MgCl_2_) containing protease inhibitors. Using bovine serum albumin as standard, we determined the total protein content by the Bicinchoninic acid method. We separated thirty micrograms of protein from cell lysates by SDS-PAGE (10% bis-polyacrylamide) and transferred them to a nitrocellulose membrane. We incubated the membrane with an anti-Bcl-2 (sc-7382, Santa Cruz Biotechnology) or anti-Caspase-3 (sc-56053, Santa Cruz Biotechnology) antibodies followed by an anti-mouse secondary antibody conjugated to alkaline phosphatase (AP) (A3562, SIGMA-Aldrich). We developed the immunoblots with BCIP/NBT color development substrate and normalized protein levels to Ponceau Red staining. We measured the intensity of bands by densitometry using ImageJ software.

### Statistical analysis

We tested ten extracts in at least three replicates in three or four independent experiments. We presented all data as mean ± standard error of the mean (SEM). We used GraphPad Prism software to perform the statistical analysis, and we considered significant differences compared to the control at a *P*-value of 0.05 or less.

## Results

### PFE induced several morphological changes in neuroblastoma cells

Cytotoxicity induces different events in cells that involve changes in their integrity. First, we observed the morphology of the cells under the microscope. After PFE treatment for 24 hr, the shape of the cells changed compared to the control. The neuroblastoma cells grow mostly attached with diamond-shaped and have some short neurite-like processes. However, the Santa María polyphenols decreased cell size and induced rounding and detachment of the dying cells (Figure 1).

**Figure 1.**
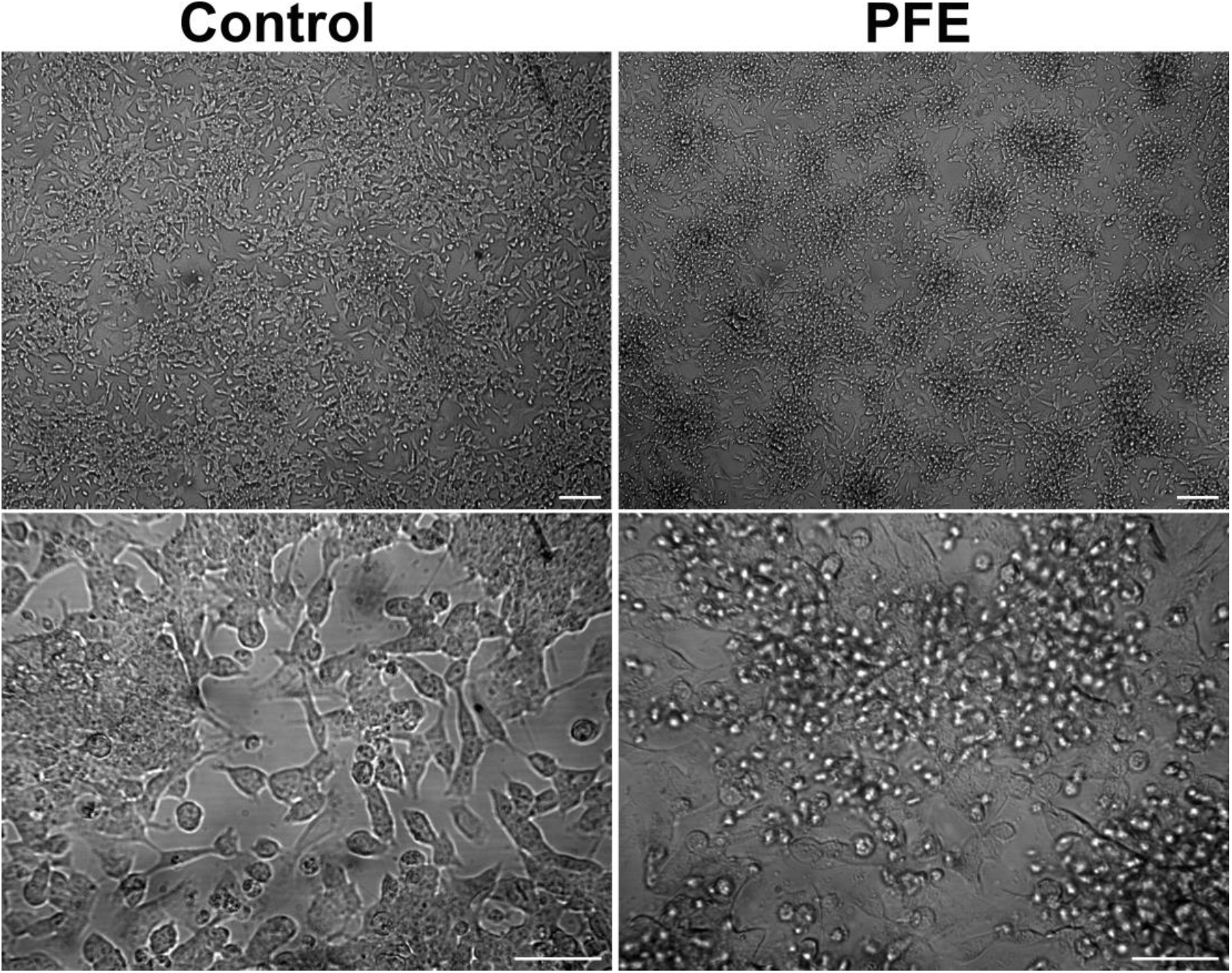
PFE induces morphological changes in neuroblastoma cells. Phase contrast microscopy images showing cell morphology from control and PFE-treated cells after 24 hr of treatment. Upper panels = lower magnification, Scale bar = 100 µm. Lower panels = higher magnification, Scale bar = 50 µm.

Alterations in nuclear morphology occur during the death of cells, although the type of changes depends on the treatment. To visualize the cells’ nuclei, we performed a DAPI staining and observed them with fluorescence microscopy. The control nuclei were large and homogenously stained with less bright blue fluorescence, while the treated nuclei were shrinkage and brighter (Figure 2A). We measured the size of the nuclei, observing a decrease of 40.66% in treated cells (*p* < 0.0001) (Figure 2B), suggesting some nuclear fragmentation. At the same time, the fluorescence intensity in these cells was higher (242.3 ± 4.89%) compared to the Control (*p* < 0.0001) (Figure 2C), indicating nuclear condensation. These results demonstrated that polyphenols induced changes in the cellular structure resulting from a controlled mechanism of death.

**Figure 2.**
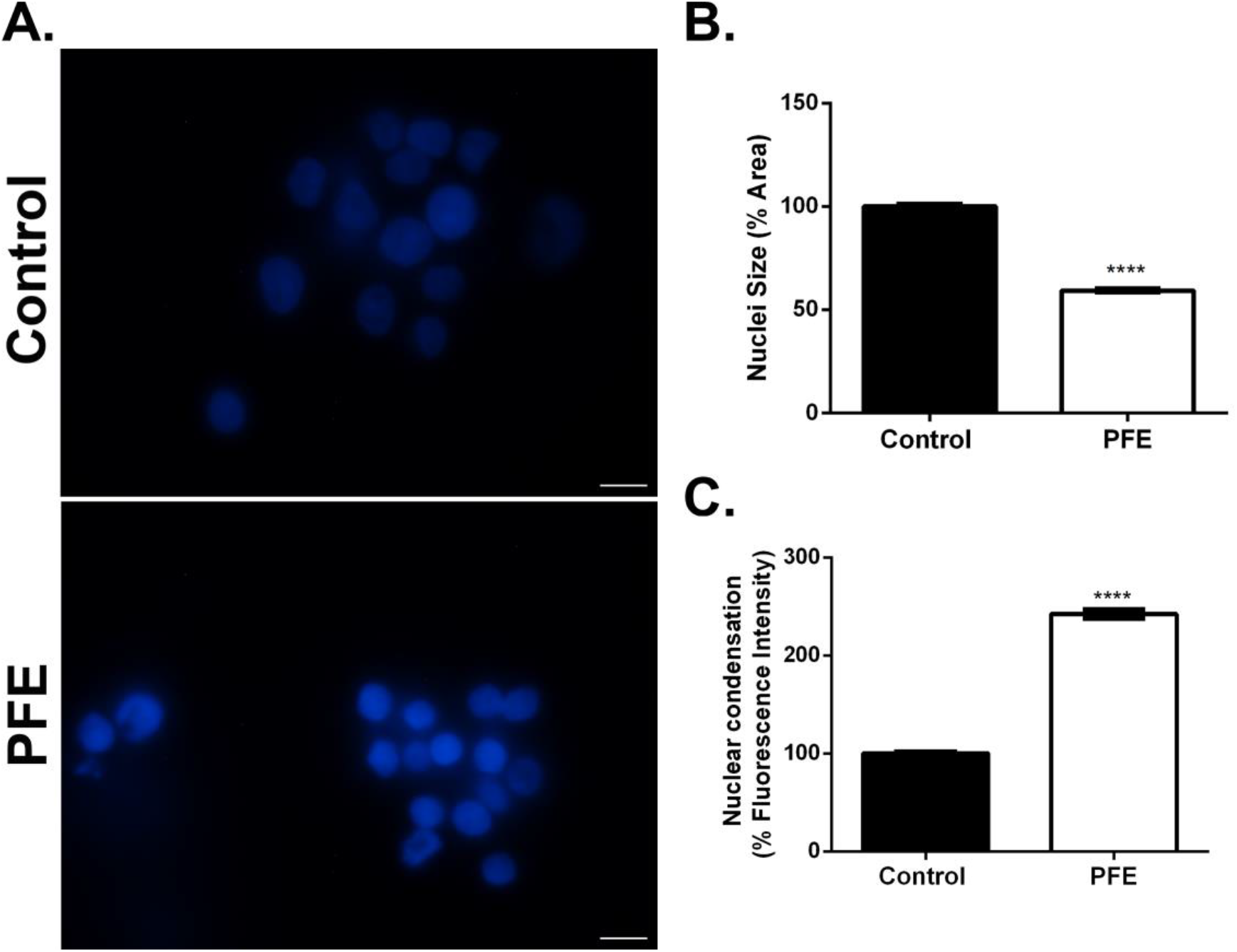
PFE induces nuclear fragmentation and condensation in neuroblastoma cells. **(A)** Epifluorescence images showing DAPI staining of nuclei in control and PFE-treated cells after 48 hr of treatment. Scale bar = 10 µm. **(B)** Measurement of the nuclei area and **(C)** quantification of mean pixel intensity. Bars represent the mean ± SEM of at least 200 cells from three independent experiments testing five extracts. **** *p* < 0.0001, Two-tailed Unpaired Student’s *t*-test.

### PFE affected the cell cycle and increased the levels of apoptotic cells

Cell proliferation involves four phases that can be affected by the treatment of some compounds. We quantified nuclei PI staining by flow cytometry to analyze the cell cycle distribution. We observed a decreased number of cells in the G1 phase after 24 hr of polyphenols treatment (data not shown). However, we found a significant cell number decrease in the G1 phase (8.46%, *p* = 0.0069) at a more extended incubation period (Figure 3A). The percentage of cells in the S and G2/M did not change. Besides, we observed an increase of cells in the sub-G1 phase (5.8%, *p* = 0.04) after the polyphenols treatment, which happens when apoptosis occurs (Figure 3B).

**Figure 3.**
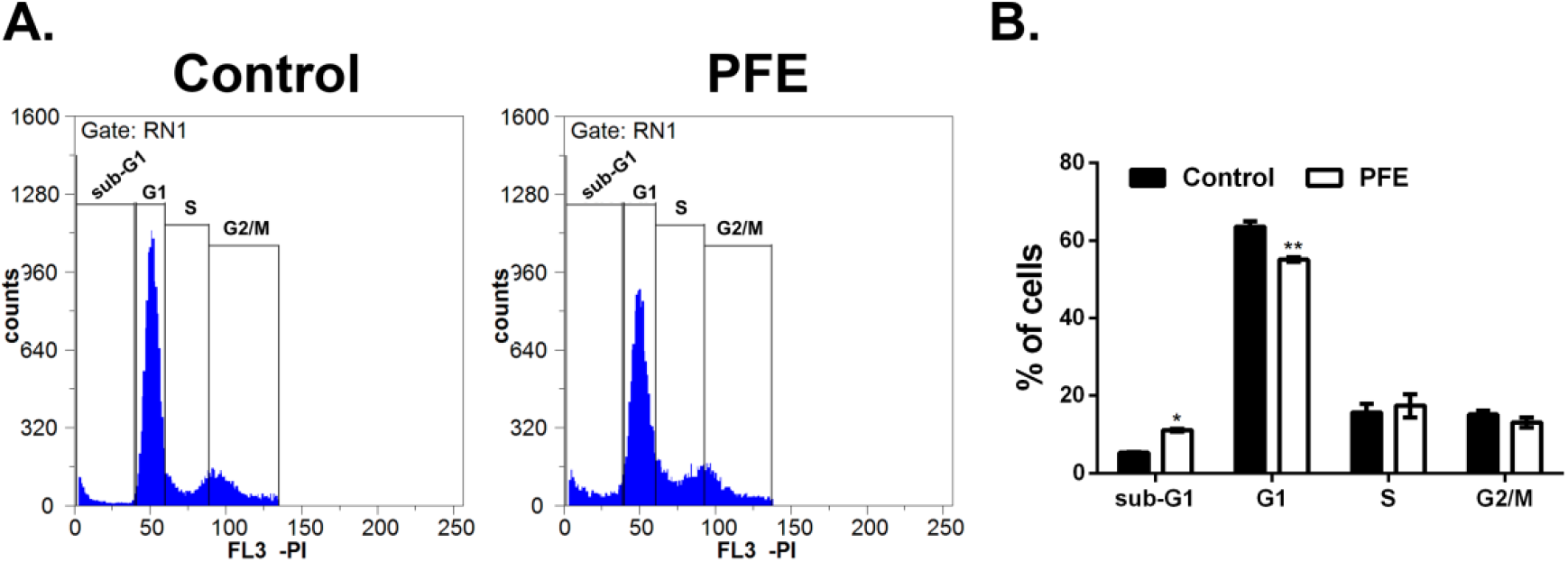
PFE slightly affected the cell cycle and increased the levels of cells in the sub-G1 phase. **(A)** Flow cytometry profiles showing the different cell cycle phases in Control and PFE-treated cells after 48 hr of treatment. RN1 = cells. **(B)** Percentage of cells in each cell cycle phase considering the control as 100%. Bars represent the mean ± SEM from three independent experiments testing two extracts. * *p* = 0.04, ** *p* = 0.0069, Two-Way ANOVA, post-Bonferroni’s multiple comparisons tests.

Apoptosis is a programmed death caused to control growth, and the cell triggers it by itself or through external signals through different pathways. To test whether the morphological changes were associated with apoptosis, we double-stained the cells with Annexin V-FITC and PI and quantified them by flow cytometry (Figure 4A). We observed a significant increase in the rate of early apoptosis (11.74%, *p* < 0.0001) (Annexin V positive but PI negative = Q4) and late apoptosis (51%, *p* < 0.0001) (PI/Annexin V positive = Q2) in PFE-treated cells compared to the control. However, we observed a minimum amount of necrosis (Annexin V negative but PI positive = Q1) (Figure 4B). In conclusion, these results suggest that Santa María polyphenols induced apoptosis as the mechanism of death.

**Figure 4.**
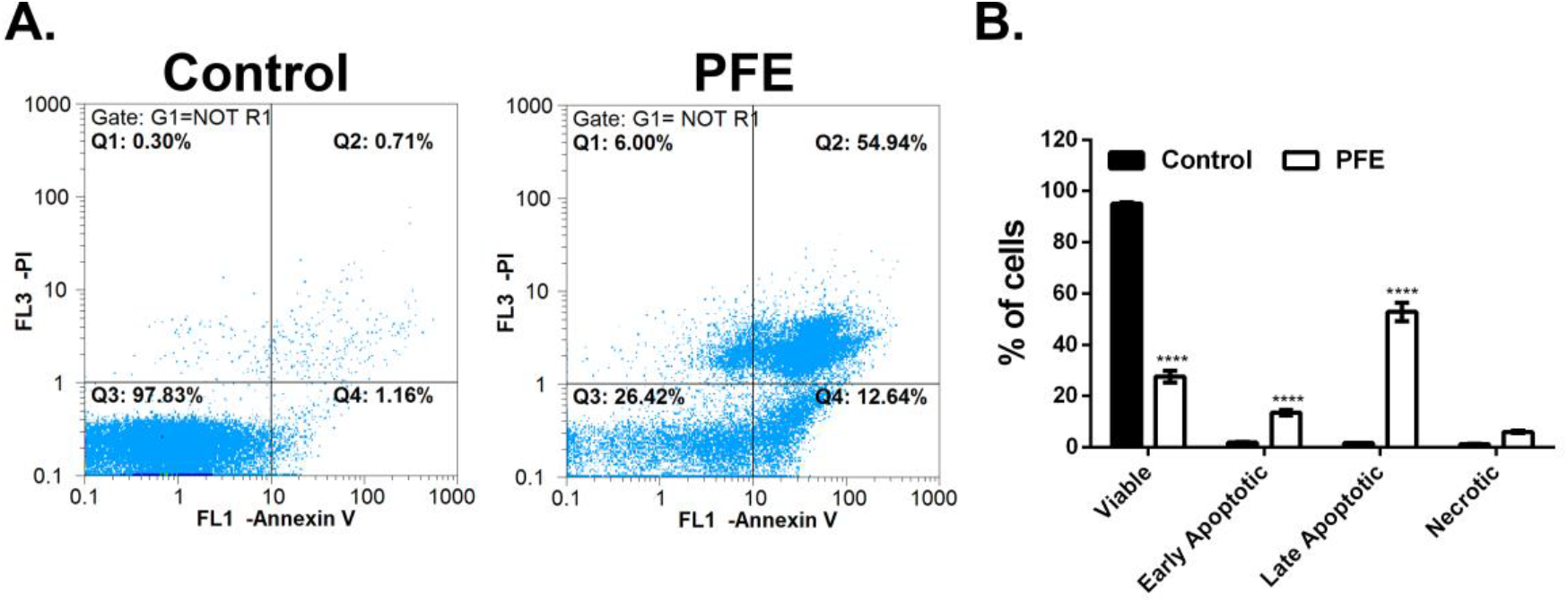
PFE increased the levels of apoptotic cells. **(A)** Flow cytometry plots showing Annexin V-FITC vs. Propidium Iodide staining in Control and PFE-treated cells after 24 hr of treatment. The four quadrants (Q1-Q4) show the distribution of cells according to their fluorescence intensity, and the software calculated the percentage of cells in each quadrant. Q1 = Necrotic, Q2 = late apoptotic, Q3 = alive, Q4 = early apoptotic. R1 = debris. **(B)** Percentage of cells in each quadrant considering the control as 100%. Bars represent the mean ± SEM from three independent experiments testing three extracts. **** *p* < 0.0001, Two-Way ANOVA, post-Bonferroni’s multiple comparisons tests.

### PFE disturbed neuroblastoma cells’ redox homeostasis and mitochondrial membrane potential

Tumor cells have high cellular metabolism to survive, generating elevated levels of reactive oxygen species compared to normal cells. To analyze whether polyphenols alter ROS levels, we treated the cells for 4 hr and measured the intracellular ROS using the probe H_2_DCFDA. Since the beginning of PFE treatment, the ROS levels decreased by around 40% (*p* < 0.0001) compared to control and were kept constant until 4 hr (Figure 5), suggesting that the polyphenols altered the redox homeostasis in neuroblastoma cells.

**Figure 5.**
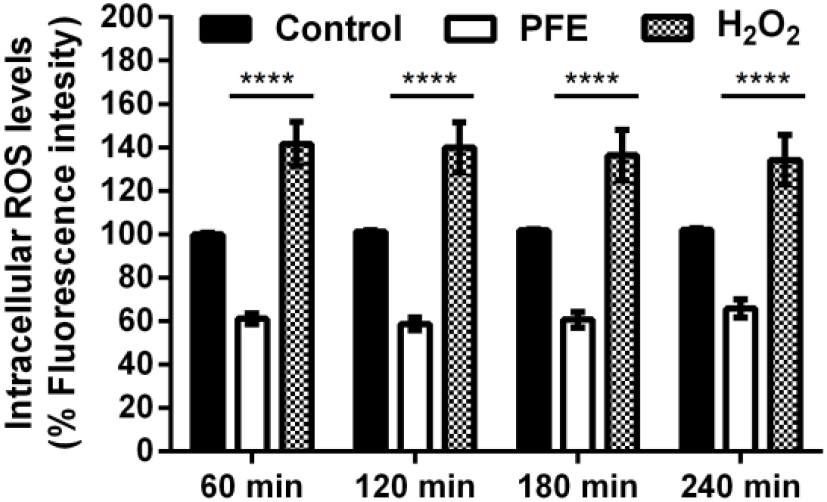
PFE disturbed neuroblastoma cells’ redox homeostasis. Determination of intracellular ROS levels measuring the fluorescence intensity of the H_2_DCFDA probe in Control and PFE-treated cells every 60 min for 4 hr. The arbitrary fluorescence units were used to calculate the percentage considering the control as 100%. Bars represent the mean ± SEM from four independent experiments testing ten extracts. **** *p* < 0.0001, Two-Way ANOVA, post-Bonferroni’s multiple comparisons tests.

The mitochondria require and maintain this homeostasis to perform metabolic functions, especially in tumor cells, where cellular metabolism is higher than usual. To test what happens in the mitochondria, we determined the potential mitochondrial membrane with Rhodamine 123. We observed a significant decrease (55%, *p* < 0.0001) in the mitochondrial membrane potential from 15 min until 1 hr of PFE treatment (Figure 6A). Then, we observed that this potential was around 58.36 ± 2.61% (*p* < 0.0001) after 3 hr of PFE treatment (Figure 6B), confirming that polyphenols induced dysfunction in the mitochondria, leading to dead cells.

**Figure 6.**
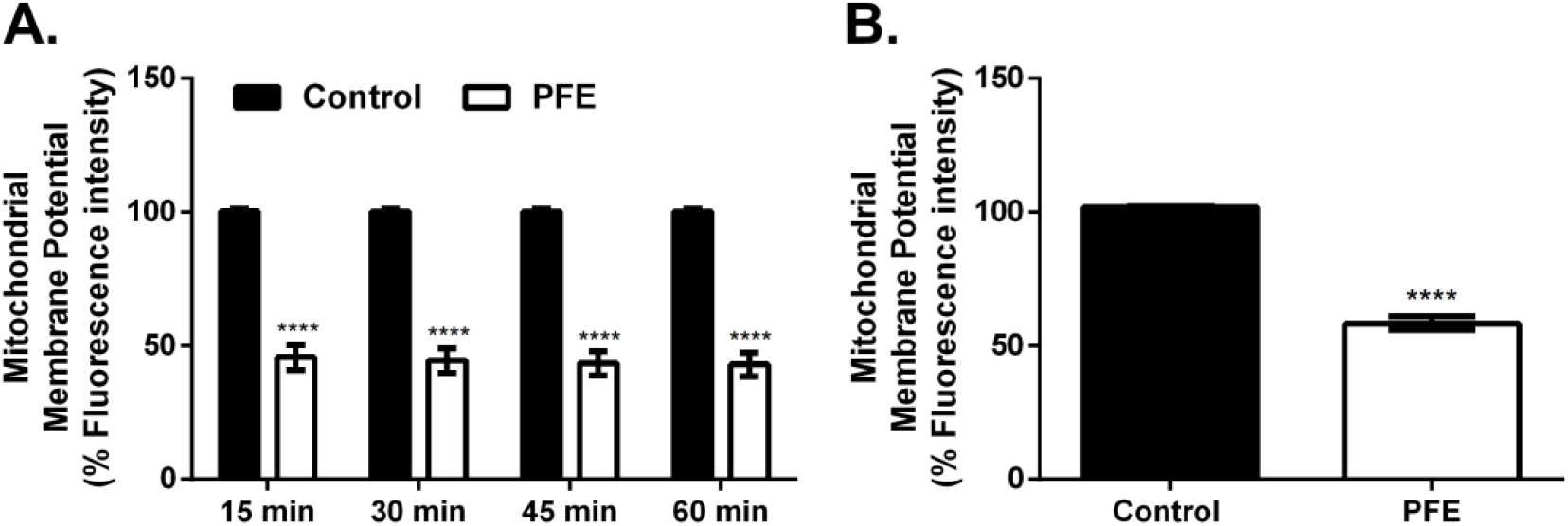
PFE induced mitochondrial dysfunction in neuroblastoma cells. **(A)** The kinetic measurement of the mitochondrial membrane potential measuring the fluorescence intensity of Rhodamine 123 in Control and PFE-treated cells every 15 min for 1hr. The arbitrary fluorescence units were used to calculate the percentage considering the control as 100%. Bars represent the mean ± SEM from four independent experiments testing one extract. **** *p* < 0.0001, Two-Way ANOVA, post-Bonferroni’s multiple comparisons test. **(B)** Determination of the mitochondrial membrane potential in Control and PFE-treated cells for 3 hr. The arbitrary fluorescence units were used to calculate the percentage considering the control as 100%. Bars represent the mean ± SEM from four independent experiments testing five extracts. **** *p* < 0.0001, Two-tailed Unpaired Student’s *t*-test.

### Bcl-2 is not involved in the PFE-induced death

In particular, the intrinsic apoptotic pathway has the mitochondria as the principal effector, where the anti-apoptotic Bcl-2 protein controls the mitochondrial membrane integrity and the cytochrome C pathway. To evaluate whether Bcl-2 was involved in the PFE apoptotic effect, we determined Bcl-2 levels by Western blotting (Figure 7A). We did not obverse changes in Bcl-2 protein levels after PFE treatment (*p* = 0.3687) (Figure 7B), indicating that this protein is not involved in the mechanism of death.

**Figure 7.**
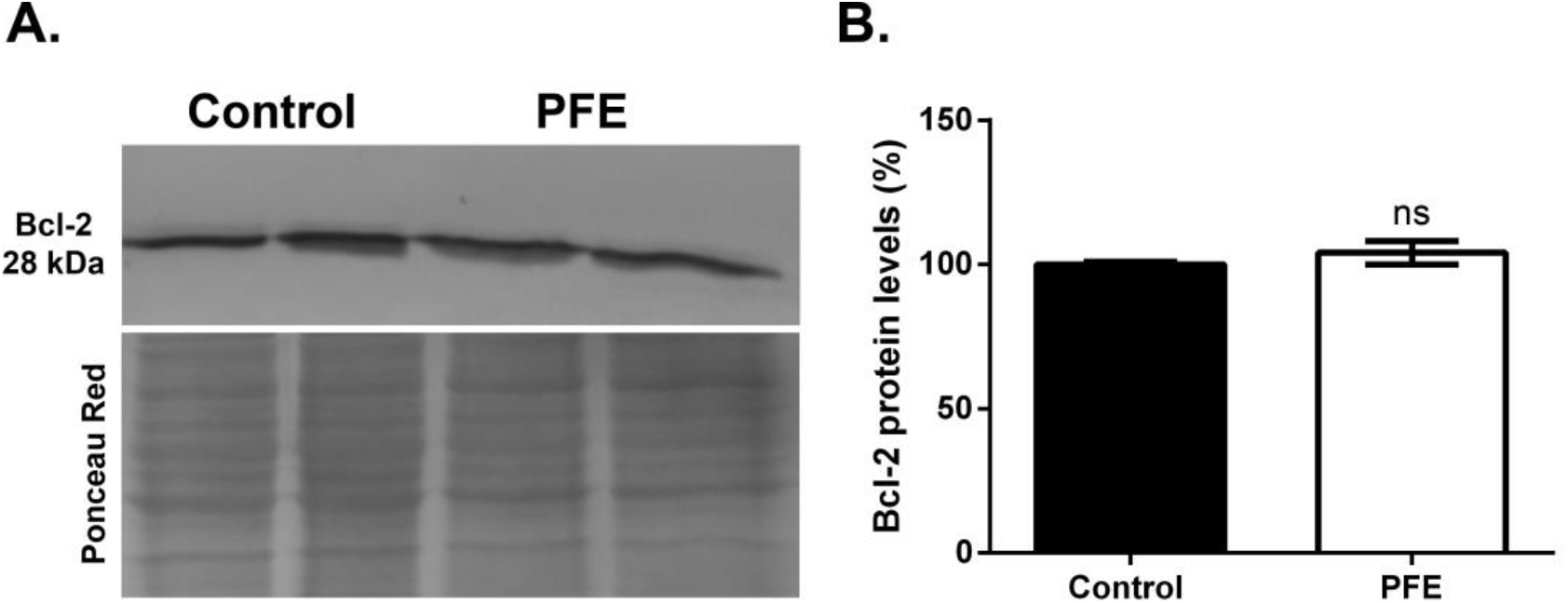
Bcl-2 is not involved in the PFE-induced death. **(A)** Immunoblot showing the Bcl-2 protein levels in Control and PFE-treated cells after 24 hr of treatment. **(B)** Quantification of Bcl-2 protein level normalized to total protein content stained with Ponceau Red (lower panel). The arbitrary units of densitometry were used to calculate the percentage considering the control as 100%. Bars represent the mean ± SEM from three independent experiments testing two extracts. No significant = ns, *p* = 0.3687, Two-tailed Unpaired Student’s *t*-test.

### PFE did not induce DNA internucleosomal fragmentation and caspase-3 activation

DNA fragmentation into internucleosomal fragments is a feature of apoptosis and depends on caspase-3 activation. We analyzed the genomic DNA into an agarose gel by electrophoresis to determine whether polyphenols induce DNA fragmentation. We observed a high molecular weight band corresponding to genomic DNA in Control and PFE-treated cells. We did not find 180-200 bp bands after PFE treatment, although the DNA size slightly decreased (Figure 8A). This result is consistent with the smaller nuclei observed in DAPI staining and flow cytometry analysis, indicating a breakdown of DNA into large fragments without further cleaved into smaller segments. Therefore, we determined the activation of caspase-3 by Western blotting. We detected the pro-caspase-3 band of 32 kDa in both Control and PFE-treated cells but not the bands corresponding to activated caspase-3 (19 kDa and 17 kDa) (Figure 8B). We did not obverse changes in pro-caspase-3 protein levels after PFE treatment (*p* = 0.8248) (Figure 8C), suggesting that polyphenols induced apoptosis through a caspase-independent pathway in the neuroblastoma cells.

**Figure 8.**
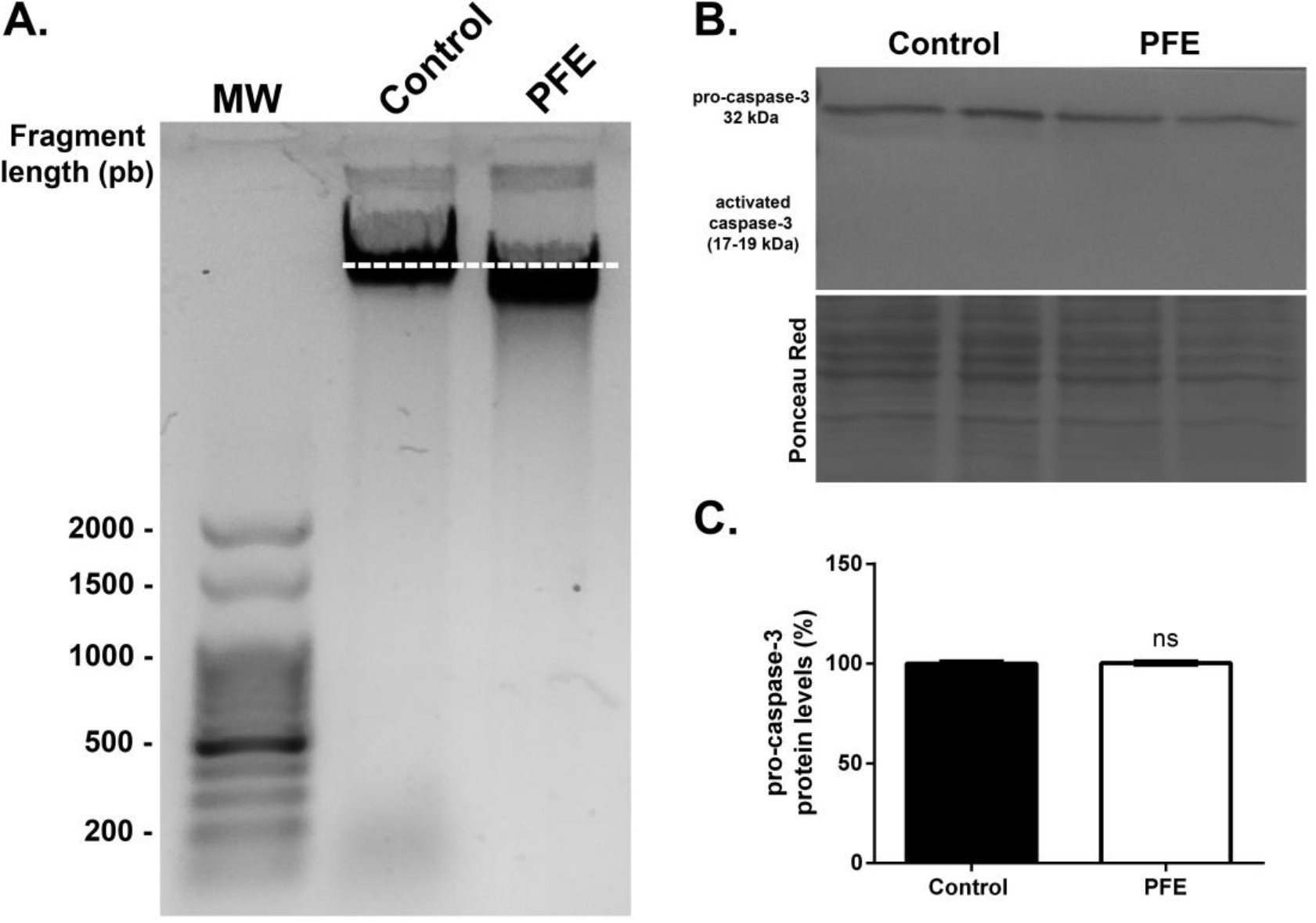
PFE did not induce DNA internucleosomal fragmentation and caspase-3 activation. **(A)** Electrophoretic analysis of the genomic DNA in Control and PFE-treated cells after 48 hr of treatment. The dashed line indicates the position of genomic DNA in control cells. MW = molecular weight marker, pb = pair bases. **(B)** Immunoblot showing pro-caspase-3 protein levels in Control and PFE-treated cells after 24 hr of treatment. **(C)** Quantification of pro-caspase-3 protein level normalized to total protein content stained with Ponceau Red (lower panel). The arbitrary units of densitometry were used to calculate the percentage considering the control as 100%. Bars represent the mean ± SEM from three independent experiments testing two extracts. No significant = ns, *p* = 0.8248, Two-tailed Unpaired Student’s *t*-test.

## Discussion

Much evidence indicates that foods rich in polyphenols could benefit human health, preventing or delaying some diseases. For this reason, identifying bioactive compounds as potential anti-tumor molecules is an exciting field of research. In the present study, we demonstrated for the first time that polyphenols from the Andean potato variety Santa María have anti-neuroblastoma properties. We showed that Santa María polyphenols extract induced morphological and physiological changes that prompted programmed cell death through the mild-oxidative state of the tumor cell, being the mitochondria the principal target.

Here, we selected a CC_50_ of 400 µg/mL of PPE to continue with the mechanistic approach. However, this concentration was higher than in our previous work with neuroblastoma cells^22^ because of changes in the cell stock and fetal bovine serum lote used for the cell culture. Also, this CC_50_ is higher if we compare it with other studies demonstrating the cytotoxic effect of polyphenols potato extracts in different tumor cell lines^13,16,19,21^. This discrepancy could be due to the compound profile of each potato variety and the differences in tumor cell lines and confluency of cells at the assay moment. Some of these studies suggested that the extracts’ anthocyanins fractions are responsible for the cytotoxic activity observed. Based on this, we could tell that anthocyanins from the Andean variety Santa María were cytotoxic for the neuroblastoma cells since anthocyanins are major flavonoid compounds in their flesh^2^. Although, we could not discard solanine and chaconine, two potato glycoalkaloids, as cytotoxic compounds. Several studies demonstrate that these two compounds induce antiproliferative activity through different pathways in several tumor cell lines^23–28^. Though, the levels of glycoalkaloids are less abundant in the flesh than in the skin of potatoes^23^, suggesting that they are not the best candidate for this effect. Another hypothesis could be that chlorogenic acid, the primary phenolic acid found in potato extract, exerts a cytotoxic effect. Some reports show that chlorogenic acid has anti-tumoral activity against colon^29,30^, osteosarcoma^31^, and breast cancer cell^28^. Although to elucidate this, we are characterizing the phenolic compounds present in the extracts by mass spectrometry and evaluating which specific fraction or combination of compounds is involved in the anti-neuroblastoma effect.

One feature of dead cells is that they suffer changes in their morphology. Here, we observed that polyphenols rounded up the cells and provoked the loss of cell-to-cell adhesion. Even though this change occurred sometime after treatment, it could be one of the initial events more than a consequence of death. According to our result, Fjaeraa and Nånberg observed that ellagic acid, a phenolic acid, affects the cell adhesion of neuroblastoma cells^32^. Recent studies show that anthocyanin cyanidin affects cell-matrix adhesion interfering in the talin-integrin complex in colon cancer cells^33^ and decreasing the focal adhesions and lamellipodial protrusions in breast cancer cells^34,35^. Another morphological change observed after treatment was nuclear condensation and fragmentation, consistent with Bontempo et al.’s observation, where potato anthocyanins induce the same effect in leukemia cells^15^. This latest observation suggested that polyphenols would trigger controlled cell death.

Since potato polyphenols can induce apoptosis *in vitro* in several cancer cells ^13–16,18,19,21^, we investigated physiological processes associated with programmed death. Similar to other pigmented varieties of potato^15,19,21^, we observed that Santa María polyphenols affected the cell cycle progression slightly and increased the percentage of cells in the sub-G1 phase. To ensure the effect of polyphenols on the cell cycle, we consider that more experiments with a shorter treatment time are required. In agreement with those reports, our results showed that Santa María polyphenols also induced apoptosis in neuroblastoma cells. In particular, the peak sub-G1 is consistent with this hypothesis, indicating a reduced DNA content. This phenomenon would result from a progressive loss of DNA caused by the leakage of fragments in apoptotic bodies detected by flow cytometry^36^. Here, we observed that the population of late apoptotic cells was higher than the early apoptotic ones after treatment, showing critical plasmatic membrane damage. However, we suggest this damage occurs because of the lack of neighboring macrophages to phagocyte the dying cells in the *in vitro* cell culture conditions^37^. Although, several studies indicate that the plasmatic membrane rupture during apoptotic death is also intrinsically programmed and is related to cell rounding, detachment, and apoptotic body formation^38^. The apoptotic action of the compounds could be by different mechanisms, depending on their nature and the cell type. Previously, we demonstrated that Santa María polyphenols have high antioxidant activity *in vitro* in a cell-free assay^2,22^, and we wondered how they interplay with intracellular ROS in neuroblastoma cells. None of the studies with potato polyphenol extract investigated this issue. Due to their metabolism, cancer cells have higher ROS levels than normal cells, keeping a particular microenvironment for growing and proliferating^4^. This feature causes tumor cells could respond differently from normal cells to the action of antioxidant compounds, such as polyphenols^6,39–41^. Our results showed that Santa María polyphenols decreased intracellular ROS levels in neuroblastoma cells. Several studies demonstrate that some polyphenols present in Santa María flesh increase ROS levels, causing toxicity in the tumor cell and leading to apoptosis^24,29,42^. Notably, they determined the ROS levels after long-time treatments, conversely to what we did in our work. Although, it is worth noting that the ROS levels seem to be slightly higher at 240 min in PFE-treated cells compared to 60 min, suggesting that ROS levels could increase in response to cell dysfunction after that time or longer. Here, we suggest that polyphenols act as antioxidant compounds rather than prooxidants, inducing changes in the homeostasis redox of tumor cells that would modify their metabolism or modulate signaling pathways^4,43,44^.

In this sense, the mitochondria are the principal players in ROS homeostasis and energy production. We showed that Santa María polyphenols affected the mitochondrial membrane potential early, suggesting that polyphenols *per se* modify the mitochondria function rapidly. Moreover, the decrease in the potential membrane was reverted when we took out the PFE after 2 hr of treatment in another cell line (data not shown), indicating that the effect would be reversible. Evidence demonstrates that polyphenols can act as protonophores and move protons across the membranes, dissipating the mitochondrial membrane potential^8^. Furthermore, this would not be the only effect that polyphenols could have on the mitochondria. Catalán et al.^7^ and Stevens et al.^45^ compiled several works demonstrating polyphenols’ action on oxidative phosphorylation, glycolysis, and mitochondrial metabolism with selectivity in tumor cells^7,45^. Therefore, we should consider Andean potato polyphenols as bioactive compounds to target mitochondria in anticancer therapy.

Based on our results, redox homeostasis and mitochondrial dysfunction were the earlier events activated in the neuroblastoma cells by the Santa María polyphenols. The mitochondrion is the principal effector in the intrinsic apoptotic pathway, and the Bcl-2 protein family regulates mitochondrial permeabilization^46^. Previous studies with potato polyphenol extracts did not measure Bcl-2 levels, while our group showed that Santa María polyphenols induced changes in Bcl-2/Bax levels in hepatocarcinoma cells^21^. In addition, some works demonstrated that solanine modifies these protein levels, indicating that Bcl-2 is an apoptotic effector in several cell types lines^26,27^. Kopustinskiene et al. reviewed data where different flavonoids, including anthocyanins, also downregulate Bcl-2 levels^47^. Likewise, chlorogenic acid induces apoptosis by reducing the anti-apoptotic protein Bcl-2 levels^48^. Instead, the Bcl-2 levels were not affected in our study, suggesting that mitochondrial membrane permeabilization occurs as an independent process, and the dysfunction in mitochondria would generate pores in the membrane and release its content.

Finally, we analyzed some hallmark markers of apoptosis, such as DNA internucleosomal fragmentation and caspase-3 activation. Similarly to our result in DAPI staining, where Santa María polyphenols induced some nuclear fragmentation, we only observed a slightly decreased in DNA size. Interestingly, we did not observe a smear pattern of genomic DNA characterized by necrosis, confirming that Santa María polyphenols induced apoptosis in the neuroblastoma cells. However, we showed a lack of caspase-3 activation after the polyphenols treatment, concluding that the death mechanism did not involve apoptotic proteases. In contrast to these results, Madiwale et al.^13^ and De Masi et al.^18^ demonstrated that polyphenols from colored potatoes activate different specific apoptotic proteases.

According to our results, we hypothesized that another effector would be involved. Reddivari et al. proved that anthocyanins-fraction from colored potatoes induced caspase-independent apoptosis through nuclear translocation of endonuclease G and apoptosis-inducing factor (AIF) in prostate cancer cells^19^. In this sense, evidence supports that a caspase-independent mechanism of apoptosis involves the participation of the mitochondrial AIF^49^. In our work, the polyphenols would act as protonophores that modify the proton gradient and disrupt the mitochondrial membrane in the tumor cell, releasing the AIF into the cytosol. Then, AIF can bind DNA and induce large-scale DNA fragmentation^50^, according to our observation of some DNA cleavage lacking internucleosomal bands.

In summary, we showed for the first time that Santa María polyphenols alter the redox homeostasis and induce mitochondrial dysfunction in the neuroblastoma cells, leading to a caspase-independent apoptotic death. These findings demonstrate that Andean potatoes are a good source of bioactive compounds with mitochondria as a target and contribute to revalorizing them as a functional food with some impact on human health. Moreover, our results put the polyphenols in value as compounds able to induce tumor cell death even in apoptotic-resistant tumors, opening new therapeutic avenues.

## Acknowledgments

We thank MSc Andrea Clausen and MSc Adriana Digilio (Germplasm Bank of INTA-EEA Balcarce, Buenos Aires, Argentina) and Lic. Patricia Suarez for providing Andean potatoes. The human neuroblastoma cells SH-SY5Y were a gift from Dr. María Elena Avale (Instituto de Investigaciones en Ingeniería Genética y Biología Molecular Dr. Héctor N. Torres, INGEBI-CONICET, Argentina). We thank Lic. Viviana Daniel (Instituto de Investigaciones Biológicas, IIB-UNMdP-CONICET) for a technical assistant with flow cytometry. Funding from Agencia Nacional de Promoción y Tecnología (PICT #0221), Consejo Nacional de Investigaciones Científicas y Técnicas (CONICET) (PIP #0762), and Universidad Nacional de Mar del Plata (UNMdP, EXA 814/17) financed this work. M.X.S. and A.B.A. are researchers from CONICET.

## Authors contribution

**María Ximena Silveyra:** Conceptualization, Methodology, Investigation, Data curation and analysis, Writing - original draft, Review & editing, Funding acquisition. **Adriana B**. **Andreu:** Review & editing, Funding acquisition. Both authors approved the final version of the manuscript.

